# SourceData - a semantic platform for curating and searching figures

**DOI:** 10.1101/058529

**Authors:** Robin Liechti, Nancy George, Sara El-Gebali, Lou Götz, Isaac Crespo, Ioannis Xenarios, Thomas Lemberger

## Abstract

Here we present SourceData (http://sourcedata.embo.org), a platform that allows researchers and publishers to share scientific figures and, when available, the underlying source data in a way that is machine-readable and findable. SourceData is unique in its focus on the core of scientific evidence—data presented in figures—and its capability to make papers searchable based on their data content and hence directly couple data to improved discoverability. SourceData aims at establishing a self-reinforcing data ‘ecosystem’ that bridges the conventional visual and narrative description of research findings with a machine-readable representation of data and hypotheses.

In molecular and cell biology, most of the data that result from hypothesis-driven research are exclusively available in the form of figures or tables in published papers. In spite of their importance for the understanding of biological processes and human disease, these data are not available in formats that would allow systematic mining and in-depth analysis ^1^. For example, a scientist interested in comparing the data from experiments testing the effect of a new drug on a specific cancer cell line currently has to rely on keyword-based searches often limited to the paper abstracts and titles. Such search results often lack specificity and the researcher then has to screen through many figures to finally be able to compare side-by-side results reported across multiple papers. With SourceData, we provide an integrated platform that makes such data-oriented searches intuitive and efficient.

The diversity of the data types and assays used across the various disciplines of molecular and cell biology makes it challenging to represent experimental results in a uniform way. To address this issue, we summarize key aspects of the experimental design first by listing the biological entities involved in an experiment and second by categorizing them into broad experimental roles. To reduce ambiguity in the nomenclature used, entities are identified by linking them to the respective entries in major biological databases. Seven types of biological entities are captured spanning several levels of biological organisations (Table 1).

**Table 1.** Resources used to link biological entities in SourceData.

| Entity type | Primary resource | Secondary resource |
| --- | --- | --- |
| **Small molecules** | ChEBI | PubChem |
| **Genes** | NCBI Entrez Gene | Rfam |
| **Proteins** | UniprotKB/SwissProt |  |
| **Subcellular structures** | Gene Ontology |  |
| **Cell lines and cell types** | Cellosaurus | Cell Ontology |
| **Tissues and organs** | Uberon |  |
| **Species** | NCBI Taxonomy |  |

Note: when there is uncertainty with regard to the exact identity of an entity, for example when the exact isoform of a protein is unknown, several external identifiers can be associated with a single entity to specifically express such residual ambiguities.

To further represent key aspects of the experimental design, entities are categorized into components that are the object of the measurement (“assayed components”) and entities, if any, that are subjected to targeted and controlled experimental interventions (“perturbations/interventions”). These two core categories are related to the concepts ‘perturbagen’ and ‘target’ in the Bioassay Ontology (BAO ^2^) and capture an important aspect of the design of experiments where multiple conditions are compared to each other in order to test whether a given perturbation (for example the presence or absence of a drug), causes a given response (for example a change in gene expression). Additional categories include ‘experimental variables’, ‘reporters’, ‘normalizing components’ and generic ‘biological components’ (see description in Supp Info).

To demonstrate the broad applicability of the system to cell and molecular biology, we developed a user-friendly web-based tool that allows rapid computer-assisted manual extraction of the metadata model described above at the level of individual figure panels based on the information provided in figure legends and on the images. Files that contain raw or minimally processed data, when available, can furthermore be uploaded and attached to the figure. As proof of principle, we have curated a compendium of over 10,000 experiments published across 23 journals. From the 621 papers processed, 368 were related to the field of autophagy, and 253 were annotated during the publication process of accepted manuscripts at four partner molecular biology journals. Out of the 12,932 experimental panels annotated, 78% included at least one ‘intervention/assayed component’ pair, supporting the broad applicability of the SourceData model. To facilitate future processing of figures of new manuscripts and integration into standard publishing workflows, a validation interface was developed to allow authors to quality check and approve the curated information.

Based on the above framework, experimentally tested hypotheses are represented as directed relationships between “interventions” and “assayed components” that co-occur in a panel. These pairwise relationships are assembled into a directed graph, where nodes are unique entities, (Figure 1a). Leveraging the connectivity of the graph, the SourceData search engine enables users to efficiently find data (linked directly to the corresponding papers) reporting the outcome of a specific experiment (for example: “search for papers reporting data on the effect of rapamycin on autophagosomes”, Figure 1b). In addition, figures can be visualized in the SmartFigure view which displays a panel in the context of related data published in other papers and allows readers to intuitively navigate from one figure to the next by following the directional relationships between entities (Figure 1c). This application is easily embeddable in webpages, for example by publishers in the online version of research articles. The structured metadata associated with each figure is accessible via a RESTful API, which allows further applications and visualizations to be built on top of the SourceData resource.

**Figure 1.**
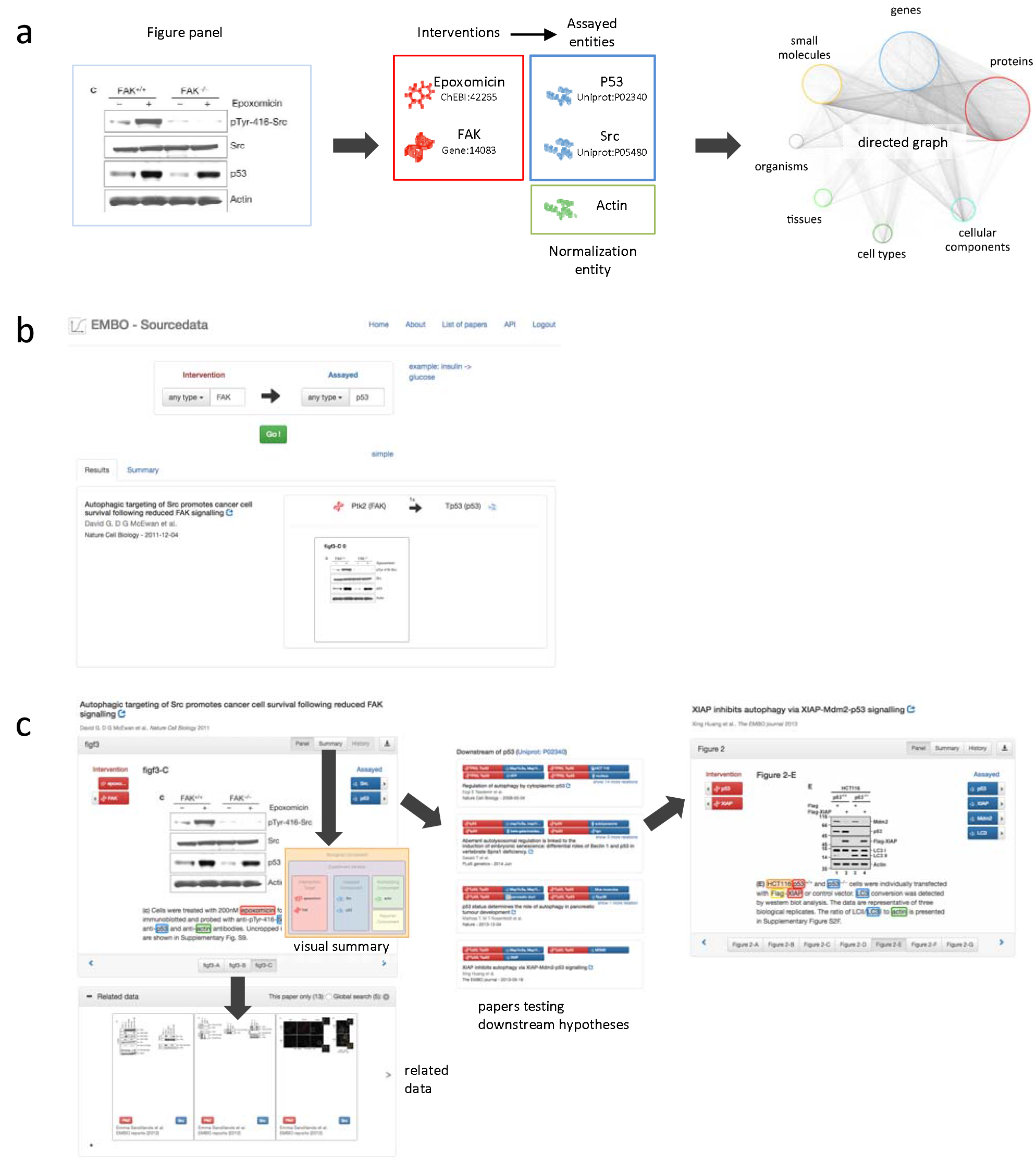
SourceData curation and navigation. (**a**) Biological entities listed in the figure (left panel) and its legend (not shown) are identified and categorized, mainly into ‘assayed entities’ and ‘interventions’, to represent the hypothesis tested in the experiment (middle panel). In the depicted example, the effect of epoxomicin and FAK are tested on p53 and Src. Such pairwise directed relationships ‘intervention/assayed entity’ between uniquely identified entities are assembled into a directed searchable graph (right panel). (**b**) The graph can be searched to find papers based on their data content. In the depicted example, the search returned a paper containing a figure showing an experiment testing the effect of FAK on p53. (**c**) The SmartFigure interface allows users to navigate through the graph from one figure to the next. It displays a figure panel in the context of related data: experiments that test similar pairs of ‘intervention/assayed entities’ (bottom panel), or experiments that test hypotheses that are either upstream or downstream (right panels) and. The view can also be toggled to a ‘visual summary’ displaying a symbolic representation of the experiments displaying entities in their respective experimental roles.

In conclusion, we provide an integrated suite of tools freely accessible to academic users, including the SourceData curation tool, search interface, SmartFigure application and public API. These resources make figures and the associated data easily searchable and will contribute to promote access to published data ^3–5^. The complete curated datasets is freely available to the community and will serve as a unique resource to benchmark the next generation of text and image mining technologies.

